# Spatial metabolomics reveals persistent localized niche-specific metabolic failure in kidneys following ischemia-reperfusion injury

**DOI:** 10.1101/2025.05.27.656326

**Authors:** Rosalie G.J. Rietjens, Benedetta Manzato, Bernard M. van den Berg, Ahmed Mahfouz, Martin Giera, Sébastien J. Dumas, Gangqi Wang, Ton J. Rabelink

## Abstract

After acute kidney injury (AKI), the persistence of failed repair proximal tubule (FR-PT) cells is postulated to hamper kidney regeneration and increase the risk of chronic kidney disease. This fibrotic shift likely depends on microenvironmental interactions, which remain largely unstudied. To investigate this, we mapped the spatial metabolic architecture of post-ischemic kidneys using an untargeted semi-quantitative spatial metabolomics (qMSI) approach, integrated with high-resolution spatial transcriptomics. Unsupervised neighborhood clustering of qMSI data revealed distinct microenvironments. Lipidome profiles identified diffusely spread areas with persistent injury markers surrounding FR-PT cells. These niches exhibited decreased linoleic acid and elevated succinic acid levels, even in epithelial cells that appeared otherwise healthy. Corresponding transcriptomic profiles confirmed downregulation of oxidative phosphorylation and fatty acid β-oxidation in these regions. Together, these findings point towards niche-specific metabolic failure and persistent mitochondrial dysfunction in areas considered healthy, underscoring the need to prioritize metabolic resuscitation to prevent long-term consequences of AKI.

## INTRODUCTION

Acute kidney injury (AKI) is becoming an increasing global health burden. The inability of the kidney to recover properly after a first insult, increases chances of AKI developing into chronic kidney disease (CKD).^1^ Upon AKI, proximal tubules can either regenerate and regain function, or fail to adapt to the injury and turn into a profibrotic phenotype, so-called failed-repair proximal tubule (FR-PT).^2,3^ This FR-PT has been focus of investigation, as it is believed that preventing this particular phenotype might aid in improving AKI outcome and preventing transition of AKI into CKD.^4^ However, focusing merely on this cell disregards the potential role of neighboring cells and the injury microenvironment on the outcome of the renal repair process.^5,6^ The effect on and the interplay with renal cells neighboring FR-PT forming injured niches have so far been relatively underrecognized, and the impact of these injured on recovery upon AKI is still largely unknown.

The rise of spatial omics techniques has paved the way to investigate the tissue microenvironment, allowing characterization of cells or agglomerates of cells within their native spatial context.^7^ While spatial transcriptomics has been most prevalent so far, other modalities are increasing in availability. From previous omics studies, it has become clear that both metabolomic^8–10^ and transcriptomic^11,12^ changes play a role in the progression of kidney injury, with metabolic alterations occurring in the acute phase upon an ischemic event, as well as in later stages with the persistence of the highly glycolytic FR-PT population. Building on these findings and technological advancements, research has recently shifted towards a multiomics approach, exploring the connection between different molecular layers to obtain a more comprehensive understanding of the processes driving kidney injury and regeneration.^13,14^

We and others have previously demonstrated that FR-PTs show metabolic impairments, both on a functional and regulatory level.^9,15,16^ The question however remains whether this impairment is only present within the affected FR-PT population, or that this metabolic effect also impedes the surrounding tissue microenvironment in its capacity to regain metabolic homeostasis upon injury. Therefore, in this work, we focused on the spatial multiomics characterization of kidney tissue two weeks after the ischemic insult to get a more comprehensive view on the interaction of FR-PT with its microenvironment.^17,18^ Spatial metabolomics revealed injury specific tissue niches, which affected the metabolic profile of PTs subpopulations. Upon niche projection, it was found that also on a transcriptomic level the niches exerted a persistent injury effect on its residing PT populations, pointing towards failure of the niche to metabolically resuscitate after an ischemic event.

## RESULTS

### Spatial multiomics approach to reveal kidney metabolic microenvironments

To characterize kidney metabolic microenvironments associated with tissue repair after injury, we induced ischemic damage to mouse kidneys with a previously described bilateral ischemic-reperfusion injury (bIRI) protocol and harvested the kidneys after two weeks for spatial molecular characterization (Fig 1). We recently established semi-quantitative mass spectrometry imaging (qMSI) which is performed using a U-^13^C-labeled yeast internal standard (IS) to allow comparison of metabolite abundances across different cell types and regions of the kidney.^18^ Based on formerly described heterogeneity of lipid species across different kidney cell types, we first performed dimensionality reduction of lipid species abundance and clustering (Supp Fig 1 – orange panel, Supp Data 1) followed by spatial mapping and overlaying with post-MSI staining using markers of known renal cell types to annotate the resulting clusters (Supp Fig 2).^9,19^ Subsequently, these lipid features were used as input for the unsupervised spatial clustering algorithm BANKSY thereby assigning individual pixels to distinct niches according to similarities of their lipid profiles and their neighborhood.^20^

**Figure 1:**
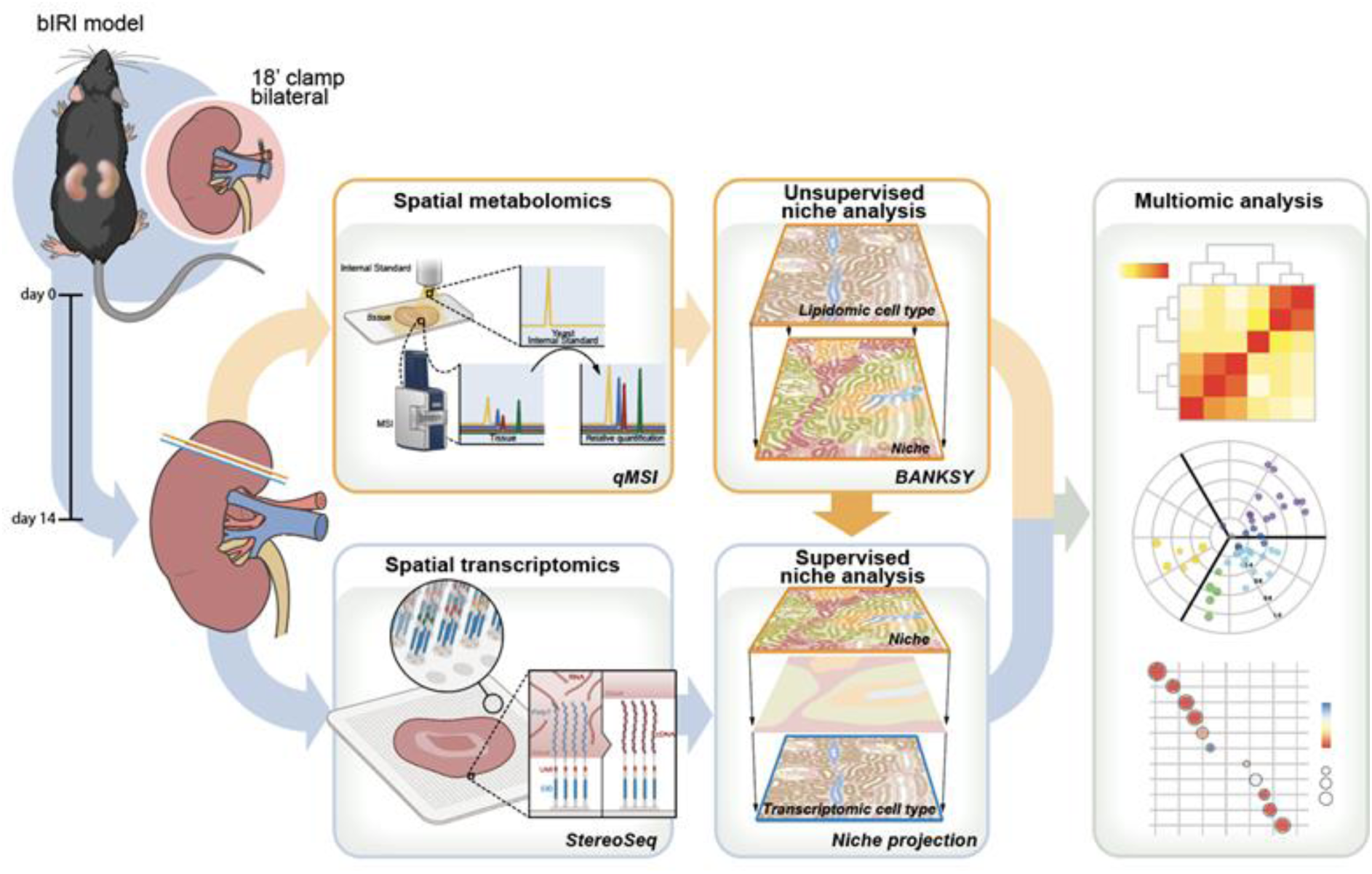
Spatial multiomics niche profiling of mouse kidneys after bIRI. Schematic representation of the experimental setup and data analysis strategy. Bilateral ischemia-reperfusion injury (bIRI) was used as model for acute kidney injury, in which the renal vasculature was bilaterally clamped for 18’ min. At day 14 post intervention, kidneys were harvested for spatial metabolomics and transcriptomics analysis. The lipidome profile was utilized for the unsupervised niche clustering algorithm BANKSY to identify areas with a persistent injury phenotype, upon which semi-quantitative MSI (qMSI) based on yeast internal standard normalization was performed. In parallel, consecutive kidney tissue sections were analyzed with STOmics Stereo-seq. The niches identified in the metabolomics modality were projected onto the transcriptomics layer to identify spots belonging to biological niches of interest in a supervised manner. Subsequently, relevant biological niches (healthy and injured) were subjected to downstream multiomics analysis.

**Figure 2:**
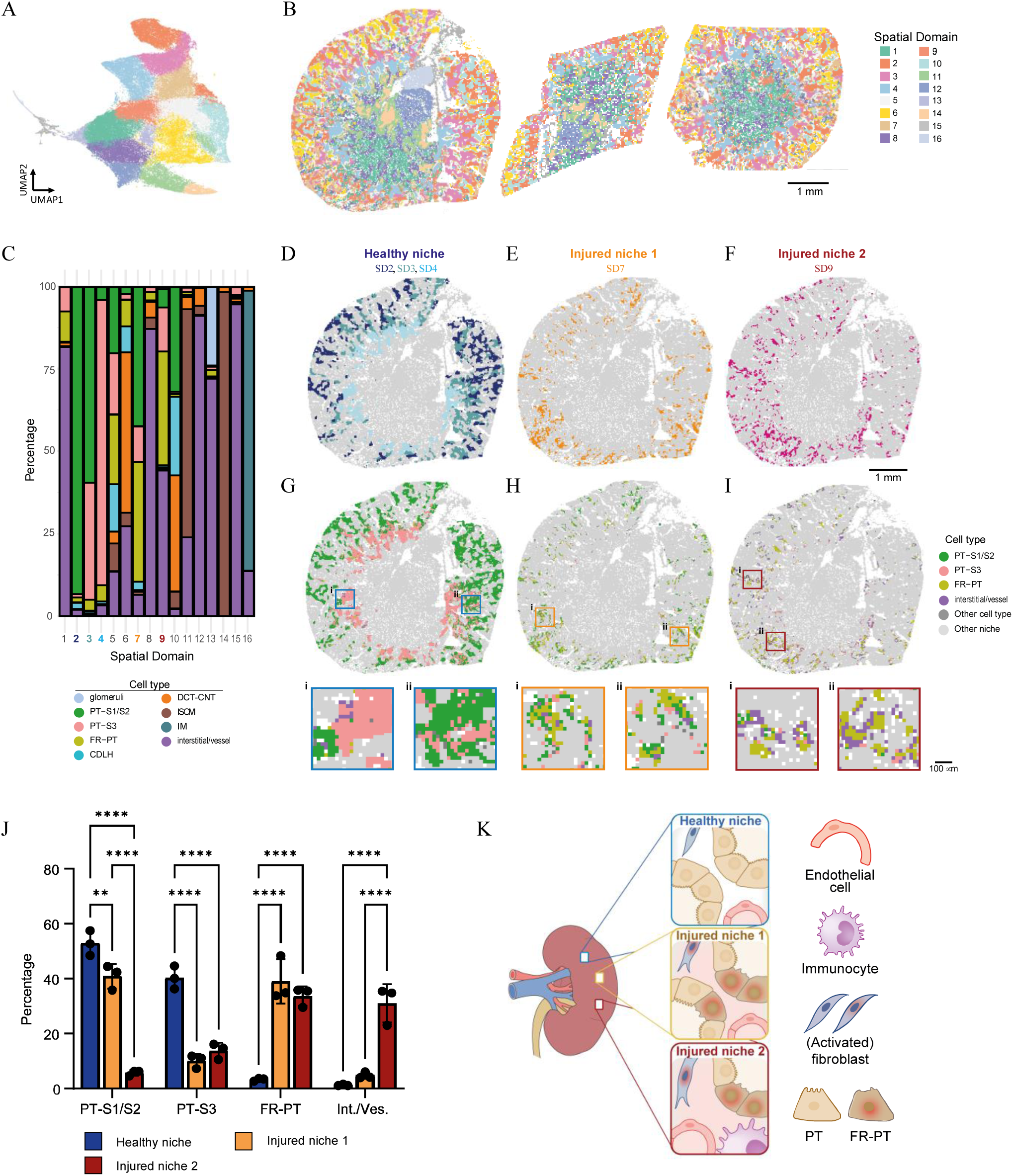
Unsupervised metabolomics niche analysis discloses distinct healthy and injured niches. Unsupervised metabolomics niche analysis was performed using the algorithm BANKSY. A: UMAP representation of the BANKSY embeddings of the bIRI mouse kidney sections (*n* = 3), colored using BANKSY (l = 1) spatial domain (SD) labels. B: Spatial maps for SD clustering for the three individual kidneys. Scale bare represents 1 mm. C: Composition of the cell type population within the SDs. Colored domain numbers (2-4, 7, 9) indicate SDs used for downstream niche analysis. D-F: Spatial maps highlighting the healthy (D), injured 1 (E) and injured 2 niches (F). Scale bar represents 1 mm. G-I: Visualization of the spatial distribution of cell types within the three niches, focusing on proximal tubule (PT) and interstitial/vessel pixel populations with more detailed views in the squared insets per niche (i, ii) displayed at the bottom panels. Scale bar of detailed views represents 100 μm. J: Bar plot of cell type counts (relative) in the healthy niche and injured niches 1 and 2. **p < 0.01, ****p < 0.0001 with 2-way ANOVA corrected with Tukey testing. K: Graphical representation of the identified healthy niche and both injured niches 1 and 2 within the bIRI kidneys. Abbreviations: PT-S1/S2, proximal tubule segment 1/2; PT-S3, proximal tubule segment 3; FR-PT, failed-repair proximal tubule; CDLH, collecting duct and loop of Henle; DCT-CNT, distal convoluted tubule and connecting tubule; ISOM, inner stripe of outer medulla; IM, inner medulla; SD, spatial domain.

In parallel, consecutive tissue sections were used for spatial enhanced resolution omics-sequencing (Stereo-seq).^17^ Similar to cell type annotation for qMSI, dimensionality reduction and clustering were applied to assign cell types based on spatial distribution and expression of canonical kidney cell type marker genes (Supp Fig 1 – blue panel, Supp Fig 3). Side-by-side comparison of the two modalities revealed a similar cell type composition in both sham and bIRI kidneys (Supp Fig 1D&I) as well as a similar spatial distribution of those cell types (Supp Fig 1 E&J), thereby supporting the validity of using the two modalities in this multiomics study.

**Figure 3:**
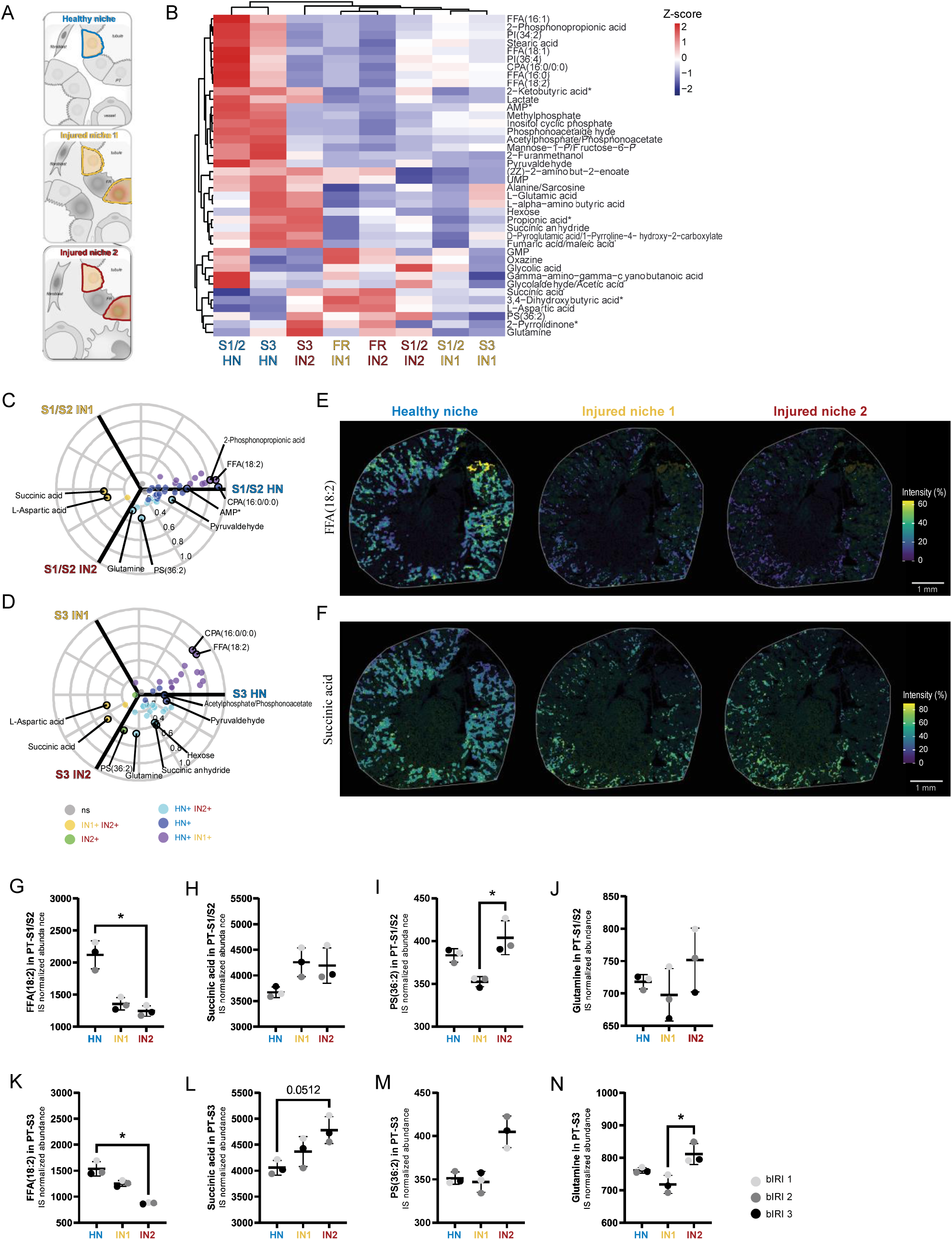
Metabolic characterization of proximal tubules in healthy and injured niches. A: Schematic of proximal tubules (PT) in different niches used for explorative downstream metabolic analysis. B: Heatmap showing niche-specific abundance of internal standard (IS) normalized metabolites in PTs as determined by semi quantitative MSI (qMSI). Color scale corresponds to z-score. C&D: Radial representation of 3D volcano analysis, comparing metabolite abundance in PT-S1/S2 (C) and PT-S3 (D) in the three niches indicated in B.^21^ Color indicates adjusted p-value < 0.05, and radial scale indicates log2 fold change. E&F: Distribution of free fatty acid (FFA(18:2) (E) and succinic acid (F) in exemplary bIRI kidney section within the three niches. Metabolite abundance is visualized as percentage of maximum value. G-J: IS normalized metabolite abundance of FFA(18:2) (G), succinic acid (H), phosphatidylserine (PS(36:2) (I) and glutamine (J) in PT-S1/S2 in the healthy and injured niches. K- N: IS normalized metabolite abundance of FFA(18:2) (K), succinic acid (L), PS(36:2) (M) and glutamine (N) in PT-S3 in the healthy and injured niches. *p < 0.05, calculated using Kruskal-Wallis test. Abbreviations: S1/S2, proximal tubule segment 1/2; S3, proximal tubule segment 3; HN, healthy niche; IN1, injured niche 1; IN2, injured niche 2.

We employed supervised niche analysis on spatial transcriptomics data with an approach we call niche projection to link the two modalities (Methods). In this approach, after manual linear alignment of the consecutive sections, cell type composition of relevant metabolic niches as identified by lipidomics- based BANSKY was used as input to identify spots in the Stereo-seq data residing in a similar niche. These regions were used for downstream multiomics analysis, focusing on characterization of the niche effect on metabolism and the metabolic transcriptome.

### Unsupervised neighborhood analysis reveals proximal tubules residing in distinct healthy and injured niches

BANKSY was applied to cluster qMSI pixels based on the lipid profile of each pixel as well as the mean and gradient of its neighborhood (k = 8). Clustering the BANKSY embeddings based on 130 lipid features (*m/z* > 400, Supp Data 1) yielded 16 distinct spatial domains (SDs), with the majority of domains having a similar distribution across the 3 bIRI kidneys (Fig 2A, B & Supp Fig 4). Of note, SDs 11, 12, 14-16 were less abundant in bIRI 2 compared to the other bIRI kidneys, which could be explained by the difference in sectioning (Supp Fig 4A,C). Some domains largely comprised of a single cell type (e.g. SD 2, 14 and 15) while the majority consisted of a combination of several cell types (e.g. SD 5, 6 and 10) indicating diffusely spread areas based on the cellular lipid profile (Fig 2C).

**Figure 4:**
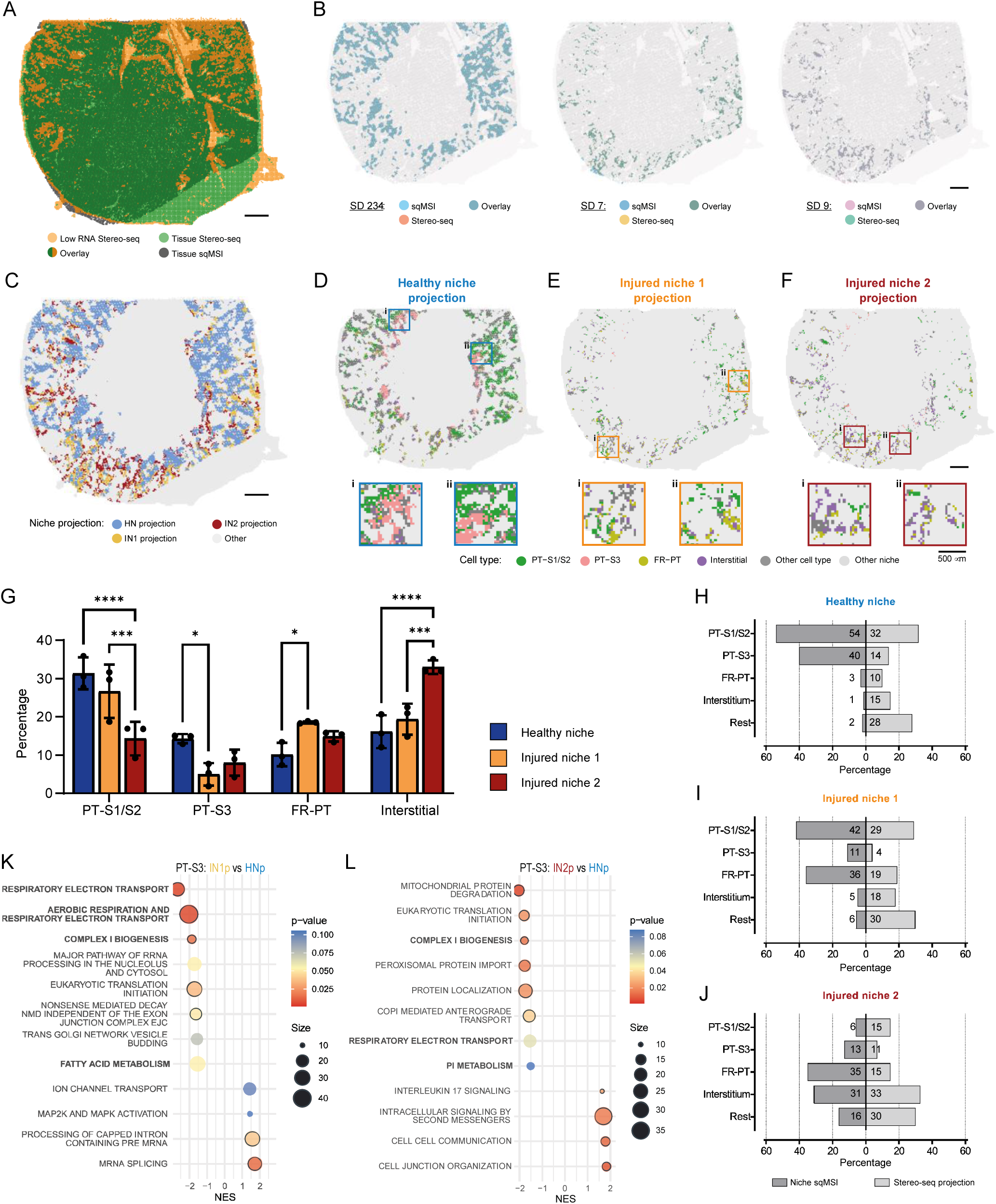
Characterization of niche projection and validation of niche-specific metabolic deregulation. A: Spatial mapping of both qMSI and Stereo-seq tissue section post linear transformation of the qMSI tissue, showing overlapping localization of gaps (qMSI) and spots containing low RNA counts (Stereo- seq). B: Visualization of spots assigned to the different spatial domains after nearest-neighbor matching of the two modalities. C: Distribution of the distinct projected healthy (HN) and injured niches (IN1 and IN2) after alignment of qMSI onto Stereo-seq. D-F: Visualization of the spatial distribution of cell types within the three niches, focusing on proximal tubule (PT) and interstitial spot populations with more detailed views of the squared insets per niche (i, ii) displayed in the bottom panels. Scale bar of detailed views represents 500 μm. G: Bar plot of cell type counts (relative) in the projected healthy niche and injured niches 1 and 2. *p < 0.05, ***p < 0.001, ****p < 0.0001 with 2- way ANOVA corrected with Tukey testing. H-J: Comparison of cell type composition (in percentage) of qMSI niches and the niches projected onto Stereo-seq. Percentages of Stereo-seq were calculated without spots annotated as ‘undetermined’. K&L: Gene set enrichment analysis of PT-S3 residing in either IN1p (K) or IN2p (L) compared to those residing in HNp. Gene sets were sorted on normalized enrichment score (NES), and the top 8 down and 4 upregulated gene sets are visualized. Nominal p- value is reported. Unless annotated differently, scale bars represent 1 mm. Abbreviations: SD, spatial domain; HN(p), healthy niche (projection); IN1(p), injured niche 1 (projection); IN2(p), injured niche 2 (projection); PT-S1/S2, proximal tubule segment 1/2; PT-S3, proximal tubule segment 3; FR-PT, failed- repair proximal tubule; NES, normalized enrichment score.

Given our primary interest in the niche effect on proximal tubules (PT), we focused on SDs predominantly composed of PTs. The lipid marker composition of PT-S1/S2 and PT-S3 was comparable between the sham and bIRI condition, therefore we hereafter regard these as healthy PT reference (Supp Fig 1K). The combination of SDs 2-3-4 was annotated as ’Healthy niche’ (HN), as these SDs contained primarily PT-S1/S2 (54%) and PT-S3 (40%) which were spatially located throughout the cortex and corticomedullary area of the kidney (Fig 2D, G, J). Furthermore, we found spatial domains (SD 7 and 9) which consisted of both a significant number of FR-PT, supplemented with regular renal cell types. This indicated a diffusely spread effect of the FR-PT to its surrounding cell types, hereafter called ‘injured niches’ (IN). Although the general spatial distribution of the two SDs throughout the kidney was similar (Fig 2E, F), their composition differed significantly (Fig 2C, H-J, Supp Table 1). For both SDs, the FR-PT contribution was similar, with 36% and 35% for SD 7 and 9 respectively. SD 7 additionally consisted of a combined healthy PT compartment of 53%, whereas interstitial/vessel pixels were predominant in the neighborhood for SD 9 (31%). This justified their consideration as distinct niches for downstream analysis, for which they are called ‘Injured niche 1’ (SD 7, IN1)) and ‘Injured niche 2’ (SD 9, IN2) (Fig 2K).

### Injured niches affect the metabolic profile of PTs

The presence of PTs in both healthy and injured niches raised the question whether their metabolic state may be influenced by the surrounding niche. We therefore investigated the effect of the neighborhood on the metabolism of PT-S1/S2, PT-S3 as well as FR-PT cells residing in either one of the three niches by comparing their metabolic profile obtained by qMSI (Fig 3A). Full processing of the qMSI data resulted in 40 annotated metabolites to be compared between the different proximal tubule cells (Supp Table 2). Both FR-PTs populations clustered together separately from the PT populations, with increased succinic and aspartic acid levels, indicating a similar metabolic profile independently of IN1 or IN2 (Fig 3B). Unlike FR-PT populations, PTs from S1/S2 and S3 clustered in a niche-dependent manner suggesting a niche-specific metabolic effect (Fig 3B). More specifically, within HN, PTs from the S1/S2 and S3 segment had a similar metabolic profile, thereby clustering separately from the PT populations residing in IN (Fig 3B). Both free fatty acids (FFA) as well as phosphatidylinositol (PI) were more abundant in PTs in the HN. Moreover, the PT populations assigned to IN displayed an altered metabolic profile compared to PTs assigned to HN, despite having the same cell type assignment according to their lipid signature (Fig 3B). PT-S3 IN2 most closely resembled tubules residing in a healthy niche, whereas PT-S3 IN1 were most dissimilar.

To quantitatively identify the metabolites associated with PTs residing in either one of three niches, we performed differential metabolite analysis (Fig 3C,D). In these plots, color indicated between which niches the metabolite is differentially abundant – e.g. yellow indicating a metabolite being up in IN compared to HN - whereas the polar angle of the metabolites indicates the extent to which a metabolite is associated with one or more niches. The largest group of altered metabolites consisted of those more abundant in PT-S1/S2 HN compared to IN (Fig 3C, HN+). For the S3 segment, it was a combination of either HN/IN1 or HN/IN2 that presented the most differentially abundant metabolites (Fig 3D, HN+/IN1+ and HN+/IN2+). Succinic acid was more abundant and linoleic acid (FFA(18:2)) was less abundant when comparing IN to HN, independently of the PT segment. Visualization of the metabolite distribution of linoleic and succinic acid validated the differential analysis, clearly showing higher intensity of linoleic acid in HN (Fig 3E) and higher intensity of succinic acid in IN (Fig 3F). For the PT-S1/S2 segment, the change of linoleic and succinic acid was similar for IN1 and IN2, whereas for PT- S3 there appeared to be an incremental change (Fig 3 G&H, K&L). Besides succinic and linoleic acid, the metabolites most differentiating between PTs in HN and IN were glutamine and phosphatidylserine (PS) 36:2. Both metabolites showed a similar trend for PT-S1/S2 and S3 for the three niches (Figure 3 I&J, M&N).

### Niche projection enables supervised transcriptomic mapping of microenvironments

Given the metabolic alterations identified, we wondered whether these changes were reflected at the transcriptional level by analyzing the consecutive tissue sections subjected to Stereo-seq. As we were mostly interested in the niche-specific effects, we sought to spatially project the lipidomic niches onto the transcriptomic data to match the metabolically interrogated qMSI pixels to their putative corresponding Stereo-seq spots (Fig 1 – Box Supervised niche analysis). We performed a linear transformation with the Stereo-seq tissue as target, where the qMSI tissue was subjected to translation, scaling and rotation to align both modalities. As the sections were consecutive, we expected gaps in the tissue which could be used as landmarks to visually validate the alignment. The gaps in qMSI were manually annotated, and for Stereo-seq spots with low RNA count (< 500) were used as pseudo annotation for gaps in the tissue. Indeed, for all three tissues the gaps in both modalities largely overlapped (Fig 4A and Supp Fig 5A). After transformation, we employed a nearest-neighbor search algorithm using the xy-coordinates of both modalities to match each qMSI pixel belonging to one of the SDs of interest with the closest Stereo-seq spot index (*k* =1). For each of the analyzed SDs, this approach yielded proper projection to the Stereo-seq tissue, for all three bIRI kidney sections analyzed (Fig 4B&C and Supp Fig B&C). The projected niches were also comparable when we enlarged the search neighborhood (*k* = 15) (Supp Fig 5DB).

We first examined the distribution and composition of the projected niches (HNp, IN1p, and IN2p) using the transcriptome-based cell type annotations (Fig 4D-G, Supp Table 3). Since the spatial location of the qMSI niches was used as input for the projection, niche distribution as well as localization of the various PT types across the corticomedullary axis remained alike in the projected niches (Fig 4D-F). HNp was characterized by an extensive healthy PT compartment (46%) (Fig 4G). IN1p and IN2p exhibited comparable levels of FR-PT (19% and 15% respectively), but could be differentiated based on either a more extensive healthy PT compartment (IN1p 32% vs IN2p 26%) or a larger contribution of interstitial cells (IN1p 18% vs IN2p 33%) (Fig 4G, Supp Table 3).

### Multiomic analysis validates niche-specific metabolic deregulation

One prerequisite for combining molecular information of the two modalities and investigating the niche-specific effects in a multiomic way, was that the qMSI and projected niches were indeed similar. Given the experimental setup – consecutive tissue sections and different molecular information for cell type annotation – we were not looking for exact correspondence, however correspondence in a broad outline of healthy and injured niches to confidently interrogate the transcriptome of PTs derived from qMSI driven projected niches. We compared the composition of the niches in both modalities, without taking along the ‘undetermined’ population in the Stereo-seq niches (Fig 4H-J). In both HN and HNp, the niche was composed for the largest part of the healthy PT compartment (94% HN and 46% HNp, Fig 4H). The FR-PT contribution of INp was slightly lower compared to IN, but was more importantly similar between IN1p and IN2p (Fig 4I-J). The healthy PT compartment of IN1 and IN1p was comparable (53% and 47% respectively), as well as the interstitial cell compartment of IN2 and IN2p (44% and 31% respectively, Fig 4I-J). The lower contribution of all PT and interstitial population is reflected in the higher number of “Rest group” cell types.

Similar to the differential analysis we performed for the metabolites, we extracted the PT populations residing in distinct projected niches to perform differential gene expression and subsequent gene set enrichment analysis (Fig 4K-L, Supp Fig 6). For PT-S3 either residing in IN1p or IN2p compared to HNp, we observed a negative enrichment score (NES) for gene sets related to respiratory metabolism, being the respiratory electron transport chain, complex I biogenesis for both IN1p and IN2p, and aerobic respiration in general for IN1p specifically. Furthermore, for IN1p, fatty acid metabolism related genes were downregulated, whereas in the IN2p population phosphatidylinositol (PI) metabolism genes displayed a negative NES. This was in correspondence with our metabolic findings, where PT-S3 in IN showed reduced levels of both FFA as well as PI (Fig 3B). On the other hand, mRNA splicing and processing as well as Mapk signaling was upregulated in the IN1p PT-S3 population (Fig 4K). For the IN2p population, it seemed that to a larger extent pathways related to (intra-)cellular communication were upregulated (Fig 4L). Taken together, these results indicate a metabolic shift towards anaerobic processes, while at the same time activating gene expression regulation and communication programs to adapt to the injured environment.

## DISCUSSION

In this study, we describe persistent localized niche-specific metabolic failure of renal epithelium after ischemic injury identified by spatial multiomics. Combining unsupervised niche analysis of the metabolomics modality with subsequent supervised niche projection onto the transcriptomics layer allowed identification and characterization of distinct healthy and injured niches, and comparison of seemingly similar healthy PT epithelial cell populations. This approach revealed a persisting localized deregulated metabolic architecture of the kidney after ischemic injury, in areas that with current approaches would be classified as normal. Spatial quantification of metabolite abundance demonstrates a similar type of metabolic failure, as has been described in the acute phase of kidney injury. Such localized areas of metabolic failure may have profound consequences for kidney function (the coutercurrent system is dependent on high energy mass transport) and cellular behaviour (persisting metabolic shifts will change the epigenome and may alter cell fate decisions)

From AKI research, it is well-recognized that accumulation of succinic acid occurs in the kidney upon an ischemic event.^22,23^ During ischemia, oxygen deprivation hampers oxidative phosphorylation leading to reversal of succinate dehydrogenase (complex II) ultimately causing succinic acid accumulation. Upon reperfusion, succinic acid is rapidly re-oxidized which generates reactive oxygen species contributing to subsequent renal injury. Attempts have been made to target and prevent succinate accumulation in order to reduce renal injury.^24–26^ Our finding of persistent high levels of succinic acid 14-days post bIRI in PTs residing in an injured microenvironment indicates that these cells might be hampered in their capacity to switch back to proper metabolic regulation after the ischemic insult. Succinic acid has been shown to fulfill several signaling roles as well; contributing to renal fibrosis by polarizing macrophages^27^ and affecting epithelial to mesenchymal transition.^28^ This combination of dysregulated metabolism and succinic acid signaling within the injured niche could lead to a maladaptive cycle of PT recovery, thereby hampering kidney regeneration.

Succinic acid is a crucial link between the TCA cycle and respiratory ETC coupled with oxidative phosphorylation.^29^ Additionally, linoleic and other fatty acids can be catabolized by beta oxidation and subsequently fuel the TCA cycle by entering as acetyl-CoA. Our results revealed that PTs from injured niches compared to those in healthy niches on the one hand had elevated levels of succinic acid, whereas on the other hand ETC and oxidative phosphorylation were downregulated. Furthermore, linoleic acid and free fatty acids were found to be less abundant in PTs in the injured niches, with concurrent alterations in regulation of fatty acid metabolism. This draws an image of metabolic dysregulation on both the entry-side as well as the metabolite flow through the TCA cycle, which will ultimately have adverse effects on the TCA cycle itself. Therefore, together the results presented in this study strongly point towards metabolic failure dysregulation in PT populations residing in injured microenvironments.

The persistent niche-specific metabolic deregulation presented in this study could carry implications for future studies into renal injury and regeneration. The FR-PT population has been a main character in the majority of recently published work, and one could think that targeting this particular cell population – either preventing its development or promoting its removal – might be the best chance to prevent AKI to CKD transition.^30^ However, recognizing the niche effect could lead to a paradigm shift where not the FR-PT population itself but its tissue microenvironment should be the target of interest. From the field of cancer^31^ and regenerative medicine^32,33^, we know how important proper microenvironmental embedding of cells is for their state and function. Three major aspects of a well-functioning, healthy microenvironment are proper vascularization, adequate lymphangiogenesis and presence of sufficient metabolic substrates. Proper vascularization and lymphatics are crucial for oxygen and nutrient delivery, as well as removal of waste products and immune cells.^34^ Persistence of an injured tissue microenvironment could lead to capillary rarefaction^35^ and disruption of the metabolic balance due to influx of immunocytes^36^, ultimately setting the resident PT population up for failure to readapt towards a healthy cell state.

In conclusion, this work highlighted the significant effect of the tissue microenvironment on its residing cell types. Without proper resuscitation of the niche, proximal tubule populations will remain hampered in their ability to regain their homeostatic cellular state, which we envision could have large implications on the longterm functional outcome for the kidney. Expanding our focus from one main character in the repair process to its theater could provide new insights useful for improving kidney performance.

## METHODS

### Mouse studies

The mouse studies and bIRI experiments were performed as previously described.^9^ In short, six 12- week-old male constitutional renin reporter (B6.Ren1cCre/TdTomato/J)^37^ mice were divided in two groups randomly (*n* = 3/group). Mice were brought under anesthesia with isoflurane and body temperature was controlled at 36.7°C. Following exposure through laparotomy, renal arteries and veins on both sides were ligated using clamps. After approximately 18 minutes, clamp removal allowed blood flow to resume (reperfusion) and the abdomen was closed. Sham control mice underwent identical surgical exposure, without ligations being performed. At day 14 post-surgery, mice were sacrificed by perfusion with cold PBS-heparin (5 UI/mL) via the left ventricle for 6 minutes at a controlled pressure of 150 mmHg before removing the kidney. Animal experiments were approved by the Ethical Committee on Animal Care and Experimentation of the Leiden University Medical Center (permit no. AVD1160020171145).

### Semi-quantitative MSI

Sample preparation, yeast internal standard (IS) and matrix application were performed as previously described.^18^ In short, kidneys were embedded in 2% carboxymethyl cellulose and sectioned into 10 μm thick sections at -20°C using a cryotome. Consecutive sections were used for the STOmics and qMSI experimental pipeline. The sections for qMSI were thaw-mounted onto an indium-tin-oxide (ITO)- coated glass slide (VisionTek Systems Ltd., Chester, UK). Slides with mounted tissues were placed in a vacuum freeze-dryer for 15 minutes and subsequently heated to ±80°C on a hot plate to denature metabolic enzymes. The yeast extracts, both unlabeled (Cambridge Isotope Laboratories, Inc. ISO1- UNL) and U-^13^C-labeled (99%, Cambridge Isotope Laboratories, Inc. ISO1) were reconstituted in 2 mL 50% methanol/deionized water. Mixtures were alternately shaken by hand and vortexed at high speed for a minimum of 2 minutes, followed by RT centrifuging for 5 minutes at 4000 rcf. Cleared yeast solutions were collected and stored at -80°C. The U-^13^C-labeled yeast solution was further diluted in methanol (1:10 v:v) for IS application. Diluted IS was sprayed on thawed ITO-slides with mounted sections, using a HTX M3+ Sprayer (HTX Technologies, USA). Subsequently with the same sprayer, N- (1-napHNl) ethylenediamine dihydrochloride (NEDC) (Sigma-Aldrich, UK) MALDI matrix^38^ solution was applied of 7 mg/mL in methanol/acetonitrile/deionized water (70:25:5 %v:v:v).

### TimsTOF MALDI-MSI

MALDI-MSI measurements were performed using a TimsTOF MALDI2 system (Bruker Daltonics GmbH, Bremen, Germany). Red phosphorus was used prior to each measurement to calibrate the instrument. Data acquisition was done in negative mode at a mass range of *m/z* 50-1000. The MALDI2 laser was set with a pulse delay time of 10 μs. All data were acquired using a beam scan area of 16 x 16 μm with a pixel size of 20 μm^2^. The laser was operated with 200 laser shots per pixel at 1 kHz.

### Immunofluorescence staining

For post-MSI IF staining used for cell type annotation, MALDI matrix and IS were washed off using ethanol and tissues were fixed using 4% paraformaldehyde for 30 minutes, then blocked with 3% normal goat serum, 2% bovine serum albumin and 0.3% Triton-X for 1h at RT. Sections were incubated overnight at 4°C with anti-CDH1 antibody (1:300, BD Biosciences, 610181) and *Lotus tetragonolobus* Lectin (LTL, 1:300, Vector Laboratories, B-1325), followed by donkey-anti-mouse AF647 (1:300, Invitrogen) and streptavidin-AF568 (1:300, Invitrogen, S11226) for 1 hour at RT. Sections were washed with PBS and incubated with DAPI ((D1306, Invitrogen) for 5 min at RT. Stained slides were embedded in ProLong Gold Antifade Mountant (Thermo Fisher Scientific, P36930). Fluorescent images were recorded using a 3D Histech Pannoramic MIDI Scanner (Sysmex, Etten-Leur, the Netherlands) or Zeiss Axio Scan Z1 Slide Scanner (Carl Zeiss AG, Oberkochen, Germany).

### MSI data preprocessing

MSI data were exported and processed in SCiLS Lab 2025a Pro (SCiLS, Bruker Daltonics). Average spectra were used in mMass to perform peak picking, with signal-to-noise ratio >10 and relative intensity threshold 0.01%. Matrix and low-quality peaks were manually excluded from the *m/z* feature list. Peak binning and transformation to a count matrix was performed with an in-house written *R* script. Count matrices were then used to create *Seurat* objects. After normalization and scaling using SCTransform, datasets were integrated using reciprocal PCA. Subsequently, UMAP dimensionality reduction and clustering with the Louvain algorithm was performed, and spatial mapping of the resulting clusters were used to establish cell type annotation.

### Identification of spatial domains

The spatially aware clustering method BANKSY^20^ was employed to perform unsupervised niche analysis. Spatial domains were identified based on 130 lipid features (*m/z* > 400, excluding low quality and matrix features). The algorithm was applied to identify spatial domains using the following parameters: *l* = 1, *k_geom* = 8, *use_agf* enabled, and cluster resolution set to 1. Domains were annotated to biological niches based on their cellular composition.

### Semi-quantitative MSI data analysis

Count matrices containing both ^13^C-labeled and unlabeled features were matched with a previously annotated yeast internal standard.^18^ Features without annotation were excluded from further analysis. For each tissue, the ^13^C-labeled and unlabeled count matrix were concatenated based on the metabolite annotation, and subsequently augmented with cell type, niche and spatial information for downstream processing. For each qMSI pixel, the normalized metabolite abundance is calculated by dividing the endogenous metabolite abundance by the abundance of the ^13^C-labeled internal standard, multiplied by the average abundance of the ^13^C-labeled IS. Metabolites with more than 20% missing values for the IS were excluded from further analysis. Succinic acid was normalized to maleic acid due to low abundance of the succinic acid IS. For the remaining metabolites, pixels with missing *m/z* values in the normalized count matrix were imputed using a smoothing approach. Specifically, the mean abundance value of the Moore neighborhood (first-order, 8 pixels) was assigned to missing-value spots. If metabolite values of all Moore neighbors were also absent, the mean of the second-order neighborhood (24 pixels) was used. In cases where no second-order neighbors contained valid values either, the overall mean abundance of the metabolite was assigned. The count matrix containing for each pixel the (imputed) normalized abundance, as well as all additional pixel information, was used for metabolic characterization of the different niches.

### STOmics Stereo-seq

Spatial transcriptomics with the STOmics Stereo-seq V.1.2 method was performed according to protocols published previously.^17^ The embedded kidneys were sectioned with the cryotome into 10 μm thick sections (consecutive section of the qMSI) and mounted to a STOmics Stereo-seq chip (1 x 1 cm) pretreated with 0.01% poly-lysine. Sectioned were incubated on a slide warmer at 37°C for 5 min, followed by methanol fixation at -20°C for 30 min. Fixed tissue sections were stained with nucleic acid dye (Qubit ssDNA assay kit) and imaged with a Motic fluorescence microscope. Subsequently, sections were permeabilized using 0.1% pepsin (Sigma, P7000) in 0.01 M HCl buffer at 37°C for 12 min followed by a washing step in 0.1x SSC buffer (Thermo Fisher, AM9770) supplemented with 0.05 U/ml RNase inhibitor (NEB, M0314L). Reverse transcription of RNA was achieved at 42°C for 3h, after which the tissue was digested with Tissue Removal Buffer at 55°C for 10 min. CDNA was released overnight at 55°C using cDNA release mix. Purification of cDNA was done using DNA Cleanup Beads (Vazyme VAHTS DNA Clean Beads, N411-02-AA) and amplified with PCR. A total of 30 ng of cDNA from the amplified product was fragmented at 55°C for 10 min and amplified once more with PCR prior to sequencing on a MGI DNBSEQ-Tx sequencer. The STOmics Analysis Workflow (SAW, https://github.com/BGIResearch/SAW) was used to perform quality control, genome alignment and quantification of gene expression of sequencing data at bin 1, and tissue-covered region selection of the data.

### Stereo-seq processing and cell type annotation

Further pre-processing and downstream analysis of the data matrices were done in R (4.3.2). For both control and IRI chips, spatial coordinates (x, y) were binned using 40 as bin size (20 µm x 20µm resolution), resulting in new spot identifiers. Unique gene and spot identifiers were indexed and mapped to integer values, which were then used to construct a sparse gene-spot matrix using the sparseMatrix function. To segment the individual tissue, a kidney segmentation mask was loaded as grayscale image for each of the tissues. This image was then converted into a numerical array and filtered to remove background spots, defining a bounding box around each tissue. Spatial transcriptomics spots corresponding to masked spots were set to zero and removed. The kidney-specific spatial transcriptomics subset was saved for downstream analysis. For each sample, Seurat objects were created, percentage of mitochondrial gene expression was quantified and data were normalized using the SCTransform v2 regularization method from the Seurat package, regressing out the percentage of mitochondrial gene expression. Subsequently, 3000 features were selected with the SelectIntegrationFeatures function and samples were pretreated with the PrepSCTIntegration function. The reciprocal PCA method, available in the Seurat package, was used for data integration using the SCT assay slot. Dimensionality reduction was performed on integrated data using PCA and subsequent UMAP embedding calculation based on the first 20 PCA dimensions. Integrated data were used for unsupervised clustering, with both resolution and algorithm set to 1. SCT assay counts were adjusted using the PrepSCTFindMarkers function, and marker genes were identified using FindAllMarkers function from Seurat. For cluster annotation, the combination was used of spatial localization, top marker genes, and canonical marker genes for kidney cell types, previously described from scRNA-seq data of similar ischemia-reperfusion injury mouse models.^4,39^ Clusters with highly similar gene signatures were merged. Clusters showing no specific cell-type gene expression were labeled as “undetermined”.

### Linear transformation for modality overlay

Linear transformation was applied in Python (v 3.12.5) to align the coordinates of the two modalities in a common coordinate framework. The Stereo-seq tissue coordinates were treated as a reference, while the qMSI tissue was used as query. For tissue IRI1, translation along the y-axis by +27,000 and x-axis by +10,000 was performed, followed by a counterclockwise rotation of 20° around the centroid. Spots with y < 25,500 were removed to account for the missing tissue in the Stereo-seq section. Then, x- and y-coordinates were rescaled based on Stereo-seq’s x and y bounds, respectively. Finally, x was translated by -400 and shrunk by 0.9. For bIRI2, translation along the y-axis by +15,300 and the x-axis by +3,150 was applied, followed by a clockwise rotation of 20°around the centroid. The x-coordinate was rescaled based on Stereo-seq’s x bounds, the y-coordinate was rescaled by 1.8 followed by a y-axis translation by +800 and x-axis by +300. For tissue IRI3, a translation along the y-axis by +13,550 and the x-axis by -100 was performed, followed by a counterclockwise rotation of 10° around the centroid. The x-coordinate was rescaled based on the Stereo-seq x bounds, while the y-axis was rescaled by 1.7. Finally, x was translated by –100. All the above-mentioned translations are in pixels.

### Niche projection

To spatially match pixels (qMSI) and spots (Stereo-seq) following tissue alignment, a nearest-neighbor search algorithm using the RANN package in R was employed. For each point in the query (qMSI) dataset, the nearest neighbor spot in the reference (Stereo-seq) dataset was identified using the nn2 function, which implements a k-d tree search algorithm. The function was executed at *k* = 1, meaning only the closest Stereo-seq spot was retrieved for each qMSI pixel based on Euclidian distance. For each qMSI pixel that had a corresponding closest Stereo-seq spot within the threshold distance (set at 50 mm), the Stereo-seq index was returned to allow identification and downstream analysis of transcriptomic niches. To identify the most abundant cell type in each neighborhood, we performed k-nearest neighbor (kNN) classification using the NearestNeighbors function from scikit-learn. We employed the Ball Tree algorithm to fit the Stereo-seq spatial coordinates as a reference and queried the 15 nearest Stereo-seq neighbors for each qMSI spot. The most prevalent Stereo-seq cell type among the neighbors was assigned to each qMSI spot, excluding ’Undetermined’ cell types.

### Gene set enrichment analysis

Niche specific PT markers were identified using the FindAllMarkers function in Seurat, with a minimum percentage of 5% across cells within a given niche and a minimum log fold change of 0.25. The resulting marker genes were used for gene set enrichment analysis (GSEA) with the fgsea package. The Reactome subset of the Canonical Pathways gene sets from the Mouse Molecular Signatures Database (MSigDB) collection was used, which was retrieved using function gmtPathways. GSEA was performed on the sorted marker gene list using the fgseaMultilevel function, with a minimum and maximum gene set size of 10 and 55 respectively. Results were visualized as enrichment score normalized to mean enrichment of random samples of the same size (NES) with nominal p-value.

### Statistics and reproducibility

All experiments and data analysis were performed on 3 animals per group. No animals were excluded from the analysis. All data are presented as mean with standard error, unless indicated otherwise. Data normality was tested using Shapiro-Wilk test. Differences between three groups were assessed by Kruskal-Wallis test, followed by pairwise comparison using Wilcoxon signed-rank test.

## ACKNOWLEDGMENTS

We gratefully acknowledge Manon Zuurmond (Department of Internal Medicine, LUMC, Leiden, the Netherlands) for making the illustrations for this article. We thank Loïs van der Pluijm, Angela Koudijs and Roel Bijkerk (Department of Nephrology, LUMC, Leiden, the Netherlands) for their technical assistance with the animal experiments. We acknowledge Marleen Jacobs and Wendy Sol for their technical support with the immunofluorescent staining. The Novo Nordisk Foundation Center for Stem Cell Medicine (reNEW) is supported by Novo Nordisk Foundation grants (NNF21CC0073729). S.D. is supported by a funding from Agence Nationale de la Recherche (ANR24-CPJ1-0181-01). T.R. is supported by the European Union through ERC grant (SPARK 101140863). Views and opinions expressed are however those of the author(s) only and do not necessarily reflect those of the European Union or the European Research Council Executive Agency. Neither the European Union nor the granting authority can be held responsible for them.

## AUTHOR CONTRIBUTIONS

Conceptualization: R.R., S.D., T.R.; Methodology: R.R., B.M., A.M., M.G., S.D., G.W.; Software: R.R., B.M., S.D., G.W.; Formal Analysis: R.R., B.M., S.D.; Investigation: R.R., B.B., G.W.; Data Curation: R.R., B.M., S.D.; Writing – Original Draft: R.R.; Writing – Review & Editing: R.R., B.M., S.D., B.B., G.W., T.R.; Visualization: R.R.; Supervision: A.M., S.D., G.W., M.G., T.R.; Project Administration: S.D., T.R.; Funding Acquisition: T.R.

