## Supplementary material for "Spatial metabolomics reveals persistent localized niche-specific metabolic failure in kidneys following ischemia-reperfusion injury": Supp Figures

### Supplemental Text and Figures

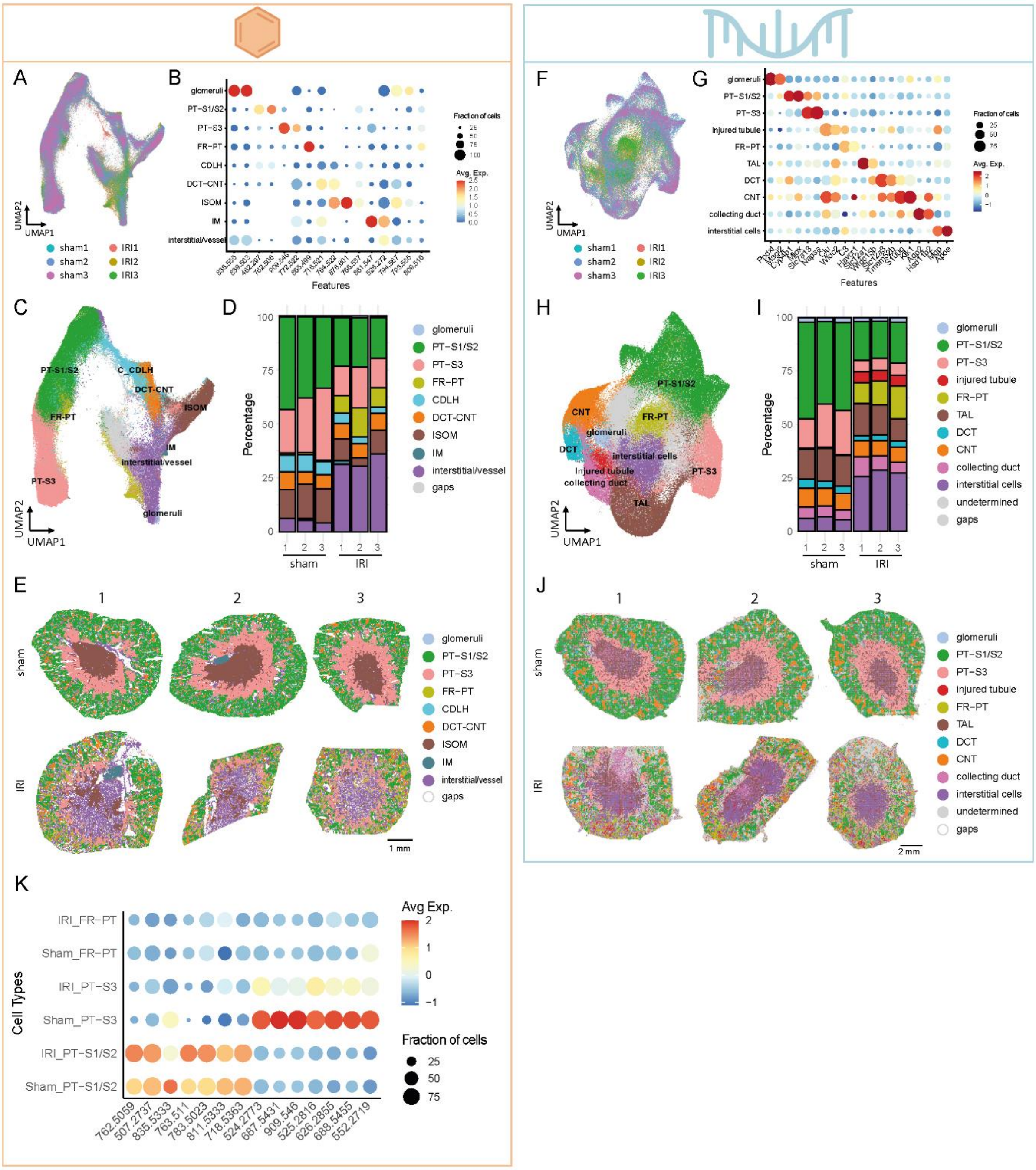

**Supplementary Figure 1: Characterization of spatial metabolomics and transcriptomics datasets.**

The orange panel (A-E, K) represents the semi quantitative MSI (spatial metabolomics); the blue panel (F-J) represents the Stereo-seq (spatial transcriptomics). A&F: UMAP visualization of analyzed kidney samples post integration (total n = 6, per group n = 3). B&G: Marker lipid (B) or gene (G) expression across the integrated sample, used for annotation of the different clusters. C&H: UMAP visualization with clusters annotated by cell type. D&I: Stacked bar plot depicting the relative contributions of each cell type across the six kidneys, revealing similar cell type composition of the samples within sham and IRI. E&J: Spatial mapping of the UMAP with pixels (qMSI) and spots (Stereo-seq) colored by cell type, showing familiar renal histology for sham and distorted histology for IRI. K: Proximal tubule lipid marker expression comparing sham to IRI proximal tubules. Abbreviations: PT-S1/S2, proximal tubule segment 1/2; PT-S3, proximal tubule segment 3; FR-PT, failed-repair proximal tubule; CDLH, collecting duct and loop of Henle; DCT-CNT, distal convoluted tubule and connecting tubule; ISOM, inner stripe of outer medulla; IM, inner medulla; TAL, thick ascending limb; IRI, bilateral ischemia reperfusion injury.

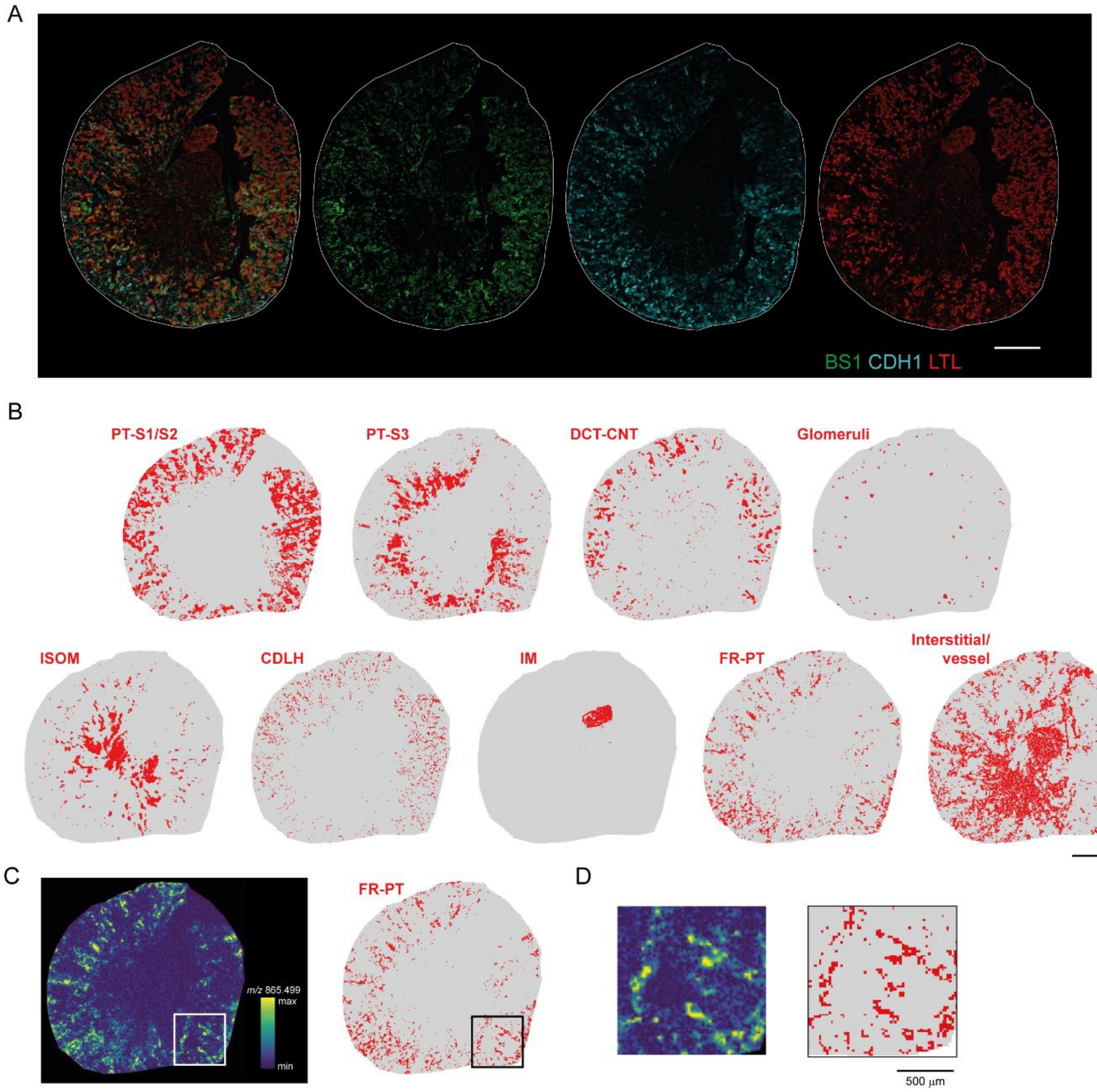


**Supplementary Figure 2: Immunofluorescence assisted cluster annotation.**

Combining immunofluorescent renal marker expression with cluster distribution and lipid marker expression allows for lipidomic cell type annotation. A: Representative immunofluorescent staining for BS1-lectin (BS1, green), E-cadherin (CDH1, blue), and *Lotus Tetragonolobus* Lectin (LTL, red). B: Distribution of various renal cell clusters on tissues as identified in Supp Fig. 1B & 1C. Scale bars represent 1 mm. C&D: Distribution of *m/z* 865.499, marker of the FR-PT cluster (Supp Fig 1B), side-by-side with the cluster distribution. Visualization of the complete kidney (C) as well as a more detailed view (D) showing correspondence of the cluster with its lipid marker.

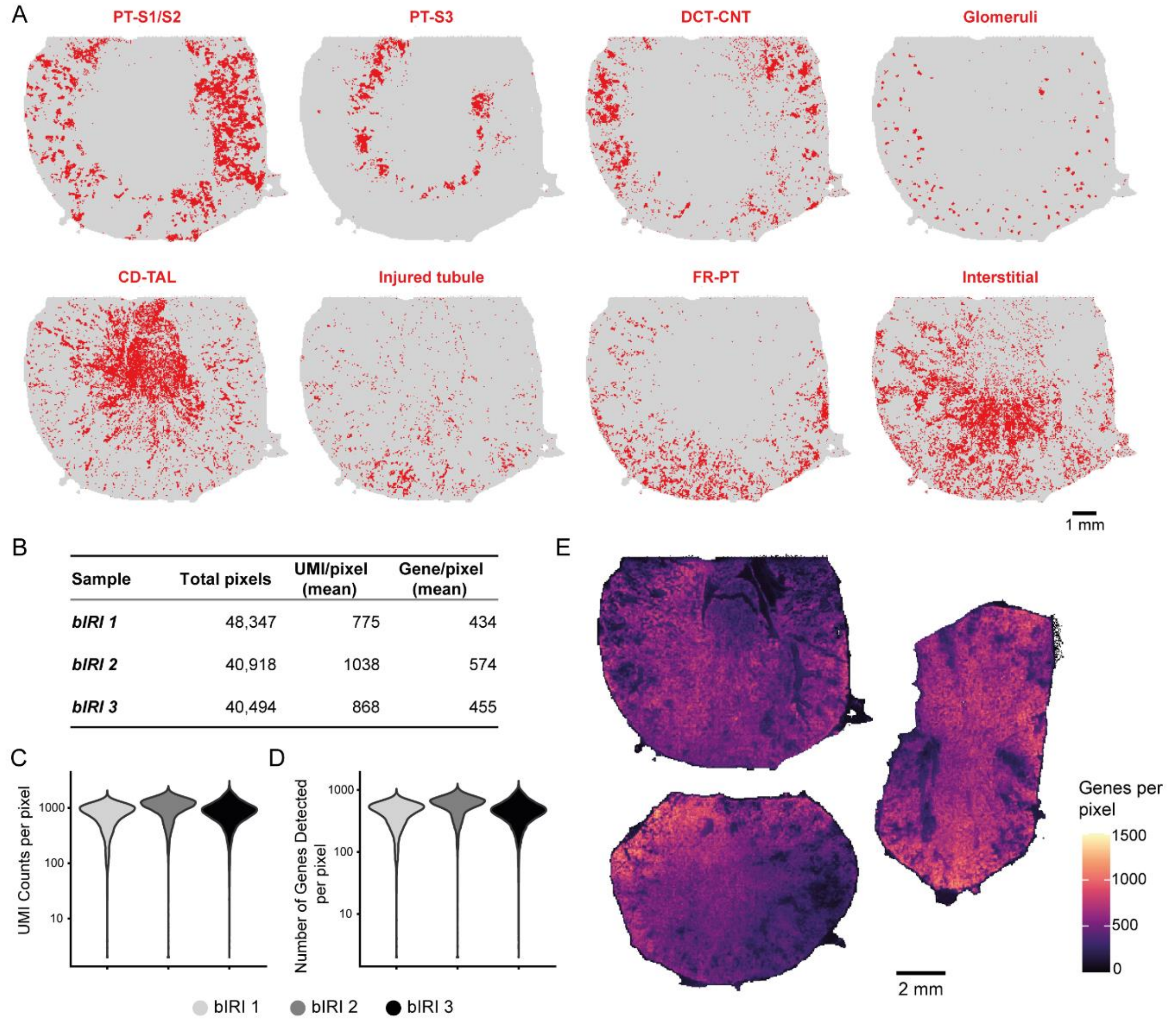


**Supplementary Figure 3: Quality control of Stereo-seq analysis.**

A: Distribution of various renal cell clusters on tissues as identified in Supp Fig. 1G & 1H. B: Overview of quality control results of the three analyzed IRI kidney sections, reporting the total spots per tissue, the mean unique molecular identifiers (UMI) per spot and mean gene count per spot. C&D: Violin plot to visualize the distribution of the UMI counts (C) and number of genes detected (D) per spot across the three IRI kidneys. E: Spatial visualization of the number of genes detected per spot. Abbreviations: PT-S1/S2, proximal tubule segment 1/2; PT-S3, proximal tubule segment 3; DCT-CNT, distal convoluted tubule and connecting tubule; CD-TAL, collecting duct and thick ascending limb; FR-PT, failed-repair proximal tubule; bIRI; bilateral ischemia reperfusion injury; UMI, unique molecular identifier.

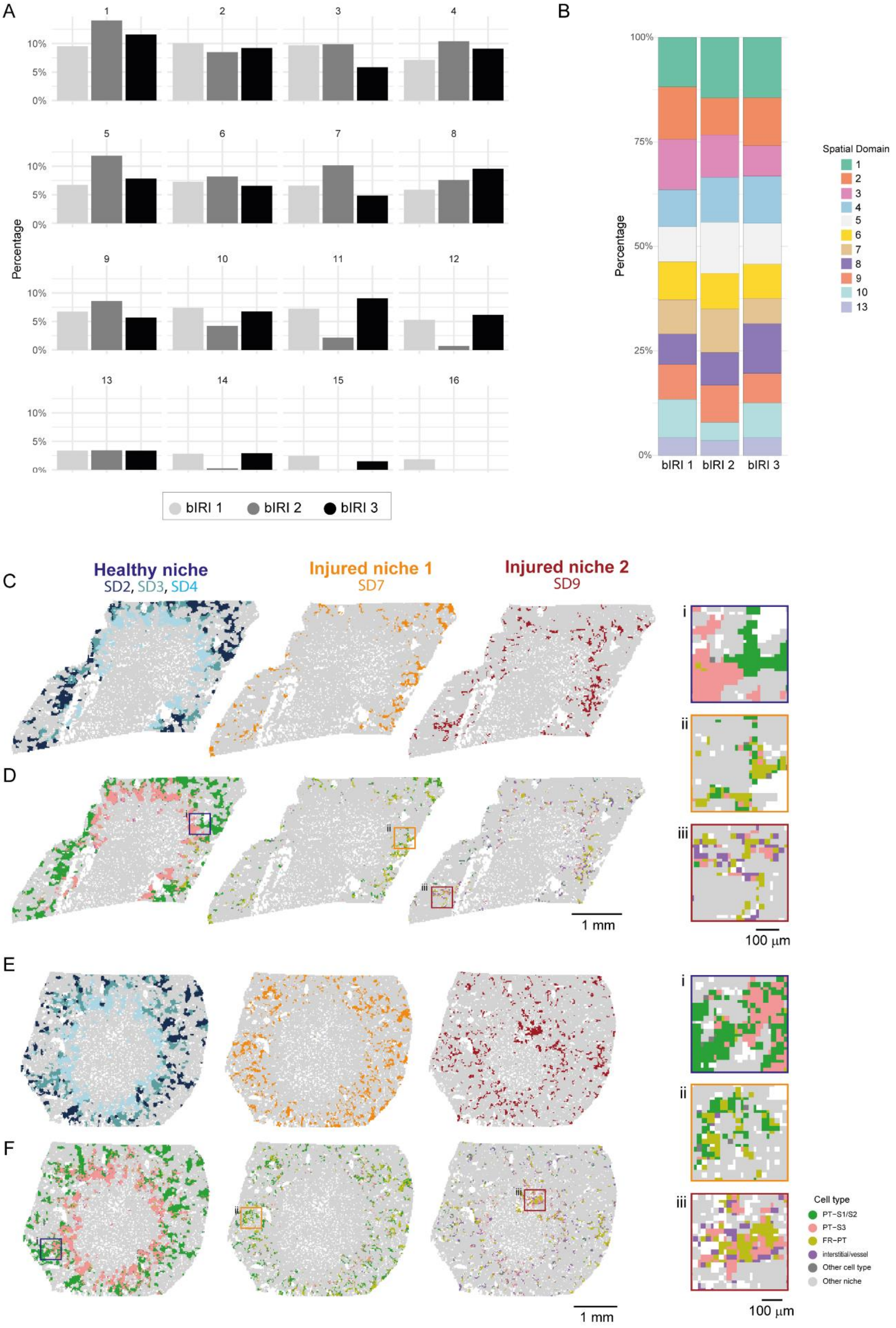

A
B
Percentage
bIRI 1
bIRI 2
bIRI 3
Spatial Domain
C
D
E
F
Healthy niche
SD2, SD3, SD4
Injured niche 1
SD7
Injured niche 2
SD9
i
ii
iii
1 mm
100 µm
Cell type
PT−S1/S2
PT−S3
FR−PT
interstitial/vessel
Other cell type
Other niche

**Supplementary Figure 4: Evaluation of BANKSY similarity across bIRI kidneys.**

A: Relative contribution of different spatial domains (SD) as identified by the BANKSY algorithm across the three bIRI kidneys. B: Stacked bar plot depicting the relative contributions of each SD across the three bIRI samples, without SD 11, 12, 14-16. C&E: Spatial maps highlighting the healthy, injured 1 and injured 2 niches in bIRI kidney 2 (C) and 3 (E). D&F: Visualization of the spatial distribution of cell types within the three niches in bIRI kidney 2 (D) and 3 (F), focusing on proximal tubule (PT) and interstitial/vessel pixel populations with more detailed views of the squared insets per niche (i, ii, iii) displayed in the right panel. Abbreviations: bIRI, bilateral ischemia reperfusion injury; SD, spatial domain; PT-S1/S2, proximal tubule segment 1/2; PT-S3, proximal tubule segment 3; FR-PT, failed-repair proximal tubule.

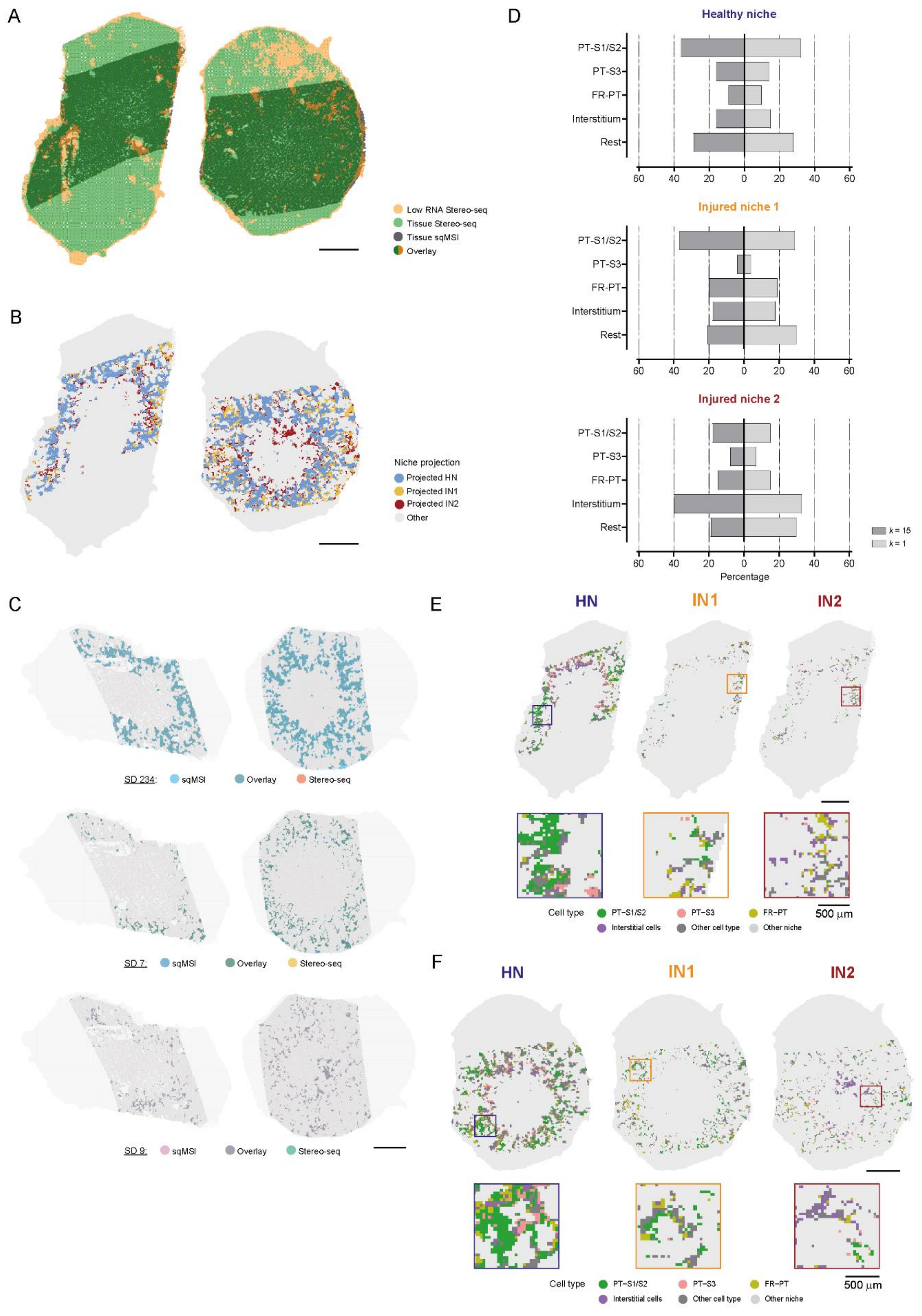

A
Low RNA Stereo-seq
Tissue Stereo-seq
Tissue sqMSI
Overlay
B
Niche projection
Projected HN
Projected IN1
Projected IN2
Other
C
SD 234: sqMSI Overlay Stereo-seq
SD 7: sqMSI Overlay Stereo-seq
SD 9: sqMSI Overlay Stereo-seq
D
Healthy niche
Injured niche 1
Injured niche 2
PT-S1/S2
PT-S3
FR-PT
Interstitium
Rest
60 40 20 0 20 40 60
Percentage
k = 15
k = 1
E
HN
IN1
IN2
Cell type
PT–S1/S2
PT–S3
FR–PT
Interstitial cells
Other cell type
Other niche
500 µm
F
HN
IN1
IN2
Cell type
PT–S1/S2
PT–S3
FR–PT
Interstitial cells
Other cell type
Other niche
500 µm

**Supplementary Figure 5: Evaluation of niche projection across bIRI kidneys.**

A: Spatial mapping of both qMSI and Stereo-seq tissue section post linear transformation of the qMSI tissue, showing overlapping localization of gaps (qMSI) and spots containing low RNA counts (Stereo-seq) for IRI kidney 2 and 3. B: Visualization of spots assigned to the different spatial domains after nearest-neighbor matching of the two modalities. C: Distribution of the distinct projected healthy (HN) and injured niches (IN1 and IN2) after alignment of qMSI onto Stereo-seq. D: Comparison of cell type composition (in percentage) of the niche projection with the closest match ($k$ = 1) and the cell type most abundant in the neighborhood ($k$ = 15). Percentages of Stereo-seq were calculated without spots annotated as 'undetermined'. E&F: Visualization of the spatial distribution of cell types within the three niches for IRI kidney 2 (E) and 3 (F), focusing on proximal tubule (PT) and interstitial spot populations with more detailed views of the squared insets per niche displayed in the bottom panels. Scale bar of detailed views represents 500 mm, all other scale bars represent 2 mm. Abbreviations: SD, spatial domain; HN(p), healthy niche (projection); IN1(p), injured niche 1 (projection); IN2(p), injured niche 2 (projection); PT-S1/S2, proximal tubule segment 1/2; PT-S3, proximal tubule segment 3; FR-PT, failed-repair proximal tubule.

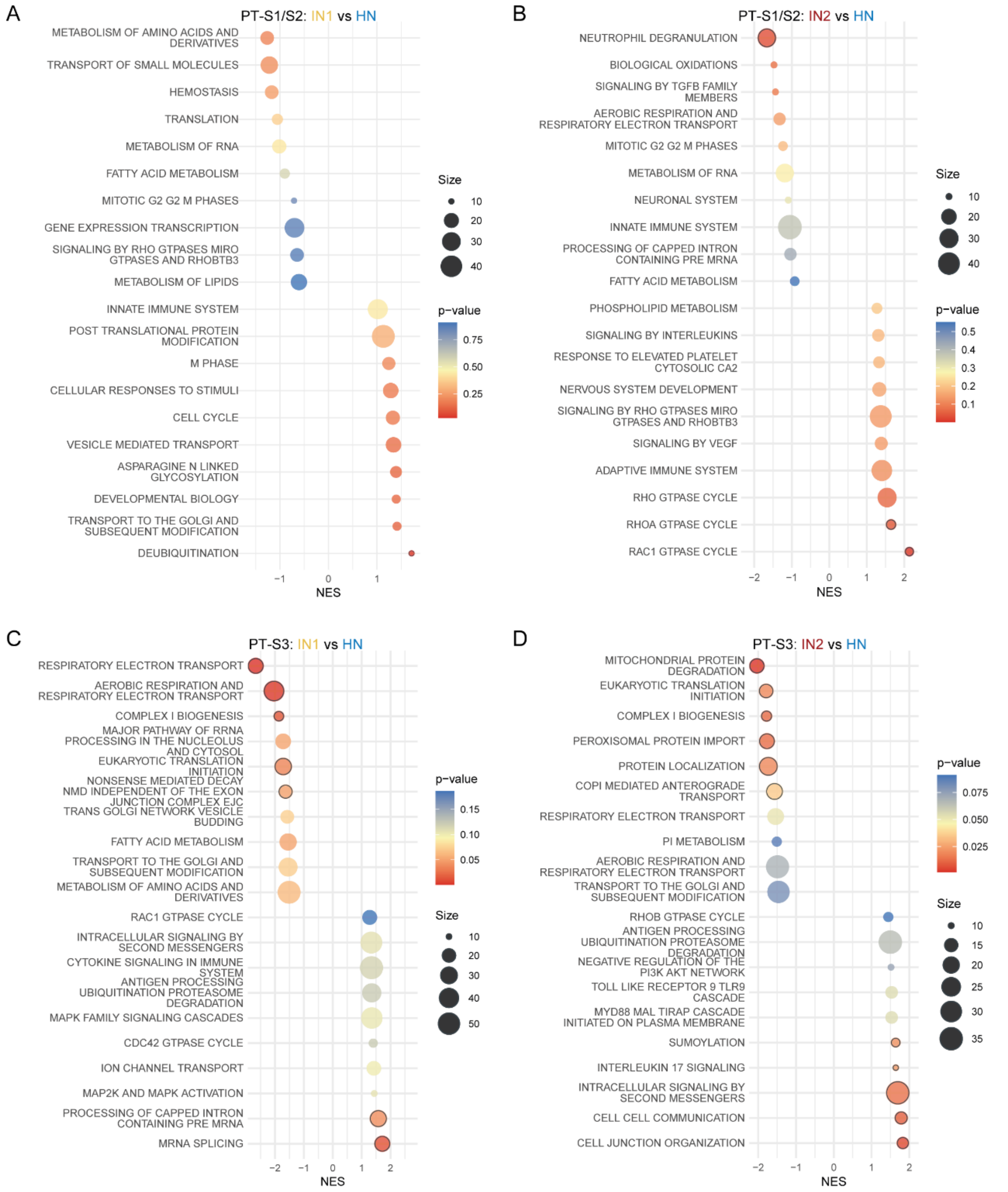


**Supplementary Figure 6: Gene set enrichment analysis of PTs from different niches.**

Gene set enrichment analysis results of PT-S1/S2 (A&B) or PT-S3 (C&D) residing in either IN1p (A/C) or IN2p (B/D) compared to those residing in HNp. Gene sets were sorted on normalized enrichment score (NES), and the top 10 down- and 10 upregulated gene sets are visualized. Nominal p-value is reported.
